# The α-Synuclein Proteostasis Network and its Translational Applications in Parkinson’s disease

**DOI:** 10.1101/2025.03.17.643594

**Authors:** Christine M Lim, Michele Vendruscolo

## Abstract

Parkinson’s disease (PD) is debilitating neurodegenerative condition that results in the loss of mobility and muscle control. A neuropathological hallmark of PD is the presence aberrant inclusions, known as Lewy pathology, of which α-synuclein (α-Syn) is a major component. The accumulation of α-Syn is a likely consequence of an age-related impairment of the proteostasis system regulating α-Syn. To investigate this phenomenon, we map the proteostasis network (PN) of α-Syn in the *Substania nigra* at the proteomic and transcriptomic levels. We then define a α-Syn proteostasis activity score (PAS) that quantifies the activity of the PN in regulating α-Syn. We thus obtain a PAS signature indicative of the disease state, as well as the age-of-death in PD patients, and the brain regional vulnerability to α-Syn aggregation. We then outline a digital twin of the α-Syn PN in the *Substantia nigra* cells by training a model on single-cell data. This digital twin is applied towards target identification for PD. In addition, we further describe the application of the PN to facilitate drug repurposing. Overall, our study highlights the implication of the α-Syn PN in PD and how simulations and measurements of its activity can help efforts in translational research for PD.

## Introduction

Parkinson’s disease (PD) is a neurodegenerative condition that progressively causes debilitating motor symptoms and cognitive impairment (1-7). Despite affecting over 6 million people globally (1-7), its molecular origins remain poorly understood, with no currently available disease-modifying treatments (8).

The aggregation of α-synuclein (α-Syn) into Lewy pathology (9, 10) is a molecular hallmark of PD (1-7). This feature that can be leveraged for the development of quantitative diagnostic methods (11, 12) and for the design of clinical trials (13). It has been proposed that α-Syn aggregates, in particular soluble oligomeric assemblies, may be neurotoxic (14-16), making α-Syn a primary target for therapeutic interventions (8, 17). The failure to maintain α-Syn in its functional state indicates that the cellular mechanisms responsible for the removal of damaged or misfolded forms of α-Syn is impaired in PD. These mechanisms are part of the protein homeostasis (proteostasis) network (PN), which regulates the behaviour of proteins in terms of their conformations, interactions, concentrations and localizations (18, 19). A defective PN has been associated with ageing and increased vulnerability to disease (20-23), suggesting that the multifactorial nature of PD may be linked with the specific impairment of PN subsystems.

In this work, we mapped the PN of α-Syn – the subsystem of the overall PN that is specifically concerned with α-Syn regulation. We then investigated the hypothesis that a disruption to the balance of the α-Syn PN to α-Syn may contribute to the accumulation of α-Syn in PD. We approached this problem by first identifying components of the α-Syn PN using functional protein interactions and transcriptomic analysis of 6 PD microarray datasets. We then validated our findings across disease progression in an independent PD dataset. Based on the α-Syn PN network, we calculated an α-Syn proteostasis activity score (PAS) that quantifies the network ability to promote or inhibit α-Syn aggregation under various conditions. Our results indicate that the PAS can identify disease states, age of death, and regional vulnerability in PD. Finally, we showed how the α-Syn PN network can be used for prioritizing targets for drug discovery and for facilitating drug repurposing.

## Results

### Building the α-Syn proteostasis network

Our initial goal was to map the α-Syn PN at the proteomic and transcriptomic levels. To achieve this result, we started by identifying the primary (first-degree) α-Syn protein interactors within the overall PN (**Methods** and **Figure 1A**), and classified them into promoters and attenuators of α-Syn aggregation based on literature **(Table S1)**. To further extend the PN of α-Syn, we identified the genes perturbed relative to *SNCA*, the gene encoding α-Syn, in PD brains **(Figure 1A)**. This analysis was based on the hypothesis that an imbalance in the relative expression of proteins involved in maintaining α-Syn proteostasis is tipped in PD, resulting in α-Syn accumulation. For this analysis, we carried out a differential expression analysis of PD and control *Substantia nigra* samples in 6 microarray datasets (see **Methods**). To account for the balance of each gene to *SNCA*, all genes were normalised to *SNCA* expression by taking log2([gene]/[*SNCA*]) for differential expression quantification. Based on our analysis, 108 genes were consistently altered (directionality) in at least half of the datasets analysed **(Table S2)**. To integrate these genes into the existing α-Syn PN network generated from protein-protein functional interactions above, we looked for functional interactions between each of the 108 genes which expression is consistently altered with respect to SNCA with the existing primary first-degree PN interactors of α-Syn. In this way, we found 21 genes that encode proteins with functional interactions with the first-degree primary α-Syn PN established earlier **(Tables S3 and S4)**. To validate the perturbation patterns of these 21 genes in PD, we analysed a separate microarray dataset consisting of *Substantia nigra* samples from PD brains of different Braak stages **(Figure S1)**, finding that the same trend of perturbation is retained. To benchmark the identification of gene markers via computing gene:*SNCA* relative expression changes, we studied the known PD genes LRRK2 and PINK1, finding that their relative expressions are indicative of brain regional vulnerability in healthy samples. In contrast, genes identified via differential expression^24^ may not always be indicative when considered in relation to α-Syn **(Figure S2)**.

**Figure 1.**
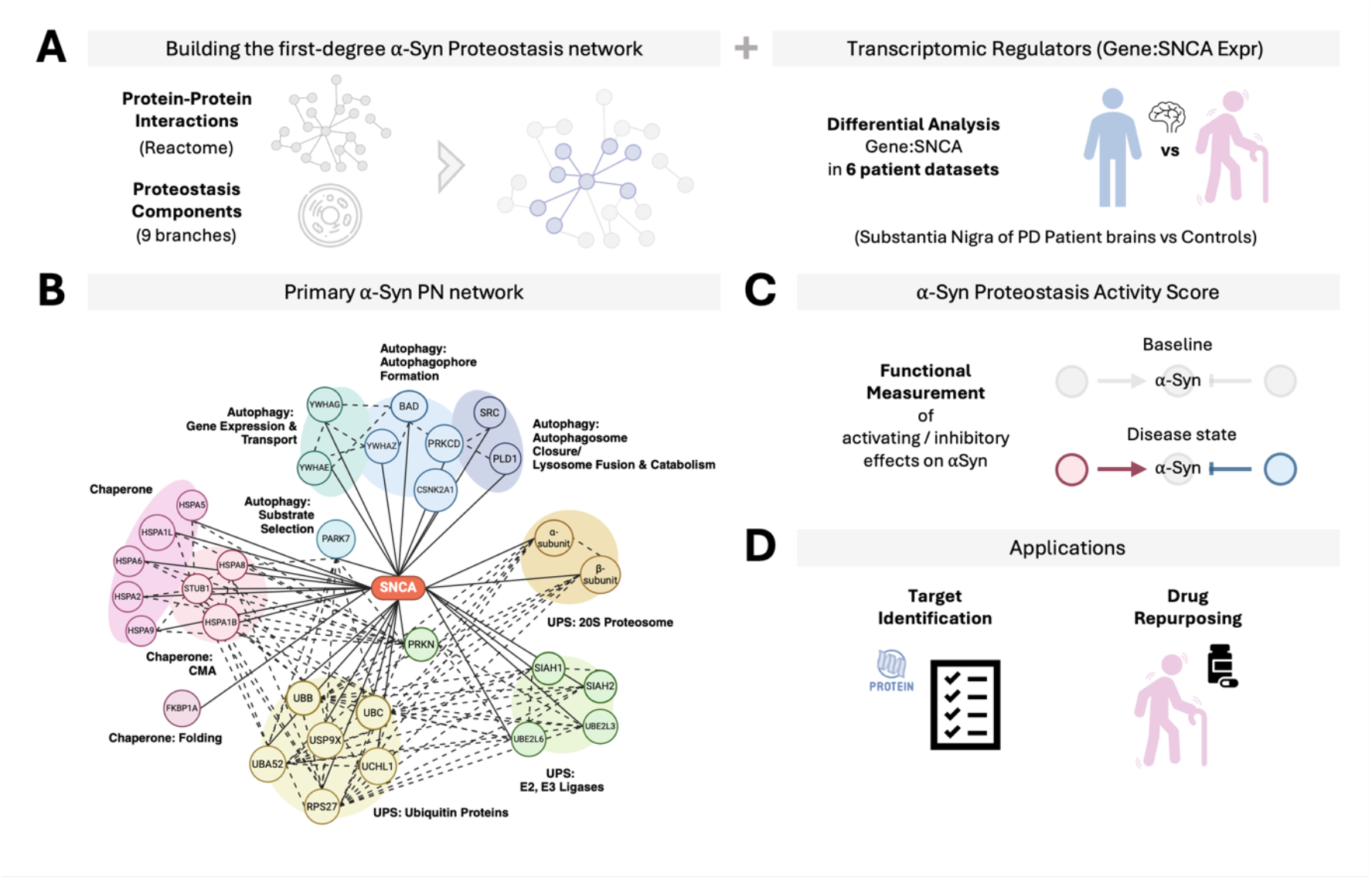
The proteostasis network of α-Syn. **(A)** To build the proteostasis network (PN), we first obtained components that are primary (first-degree) α-Syn interactors. Proteins making up the primary α-Syn PN network were then classified into promoters and attenuators of α-Syn aggregation based on the literature **(Table S1). (B)** To further extend the PN of α-Syn, genes perturbed relative to SNCA in PD brains were added to form our final α-Syn PN. The first-degree PN regulators of α-Syn are grouped into three main degradation pathways: Chaperone Mediated Autophagy (CMA), Ubiquitin-Proteosome System (UPS), and Autophagy-Lysosome Pathway (ALP). **(C)** To comprehensively quantify the functional activity of the α-Syn PN, we calculate a α-Syn proteostasis activity score (PAS) that is indicative of PD states, age of death and regional vulnerability. **(D)** We exemplify the application of our network and PAS for 2 uses: target identification and drug repurposing.

### The α-Syn proteostasis network

The α-Syn PN is presented in **Figure 1B**. The majority of the α-Syn PN is made up by molecular chaperones and by degradation systems, including Chaperone Mediated Autophagy (CMA), the Ubiquitin-Proteosome System (UPS), and the Autophagy-Lysosome Pathway (ALP). This structure of the α-Syn PN is consistent with previous reports indicating that a reduced ability to degrade damaged or misfolded α-Syn is a driver of α-Syn aggregation in PD and related synucleinopathies (24-27), and it represents a possible therapeutic target (8, 17, 28).

CMA is one of the main pathways to remove potentially cytotoxic forms of α-Syn in PD (24). In this process, α-Syn is recognized by cytosolic molecular chaperones and transported into lysosomes where it is degraded by proteases (29, 30). In familial PD, mutant forms of α-Syn are poorly degraded via CMA as mutant α-Syn binds CMA receptors resulting in a blockage of CMA degradation (24). In a similar fashion, downregulated CMA activity due to other environmental circumstances may reduce α-Syn clearance promoting buildup in sporadic cases of PD. UPS is also a well-established degradation pathway for α-Syn clearance where dysfunctional UPS may contribute to rising levels of α-synuclein in PD neurons (31). This possibility is supported by reports of reduced rates of proteasome catalytic activity (25) and lower levels of proteasome subunits in PD brains compared to healthy controls (32). Furthermore, the inhibition of the UPS was found to trigger PD neuropathology (33, 34). Given these findings, an impairment of the UPS is a likely contributing factor to α-Syn aggregation in PD.

ALP also plays an essential role in α-Syn degradation, and it is necessary for preventing PD-related α-Syn aggregation (31). Inhibition of autophagy increases α-Syn accumulation and aggregation, while its activation promotes the clearance of α-Syn inclusions (35, 36). Temporal changes in α-Syn accumulation have also been observed in accordance with changes in key ALP markers (35).

In the following sections, based on the α-Syn PN, we first generate a score to quantify the overall activity of the network on α-Syn **(Figure 1C)**, and then illustrate its use for two different applications - target prioriziation in drug discovery and drug repurposing **(Figure 1D)**.

### The a-Syn PN activity is indicative of PD states and regional vulnerability

To describe the functional biological activity of the network, we used the PAS to capture the coordinated activity of the proteins within the a-Syn PN under different states. Details on the calculation of the PAS are described in **Methods**. We found that the PAS is indicative of PD disease states – PD patients across 4 unique datasets have a higher PAS than controls (**Figure 2A**), reflecting increased activity in promoting α-Syn aggregation compared to controls. In addition, PD patients with higher PAS had a significantly younger age of death **(Figure 2B)**. In patients with PD, the peripheral nervous system (PNS) had a higher PAS than the central nervous system (CNS) (**Figure 2C**), reflecting the higher vulnerability of the vagus nerve to be compromised to α-Syn aggregation in PD progression. We also analysed two independent datasets (UKBEC (37) and GSE7307), including samples from multiple brain regions of non-disease patients, to study regional vulnerability. We found that brain regions of higher vulnerability had lower baseline levels of proteostasis activity **(Figure 2D)**.

**Figure 2:**
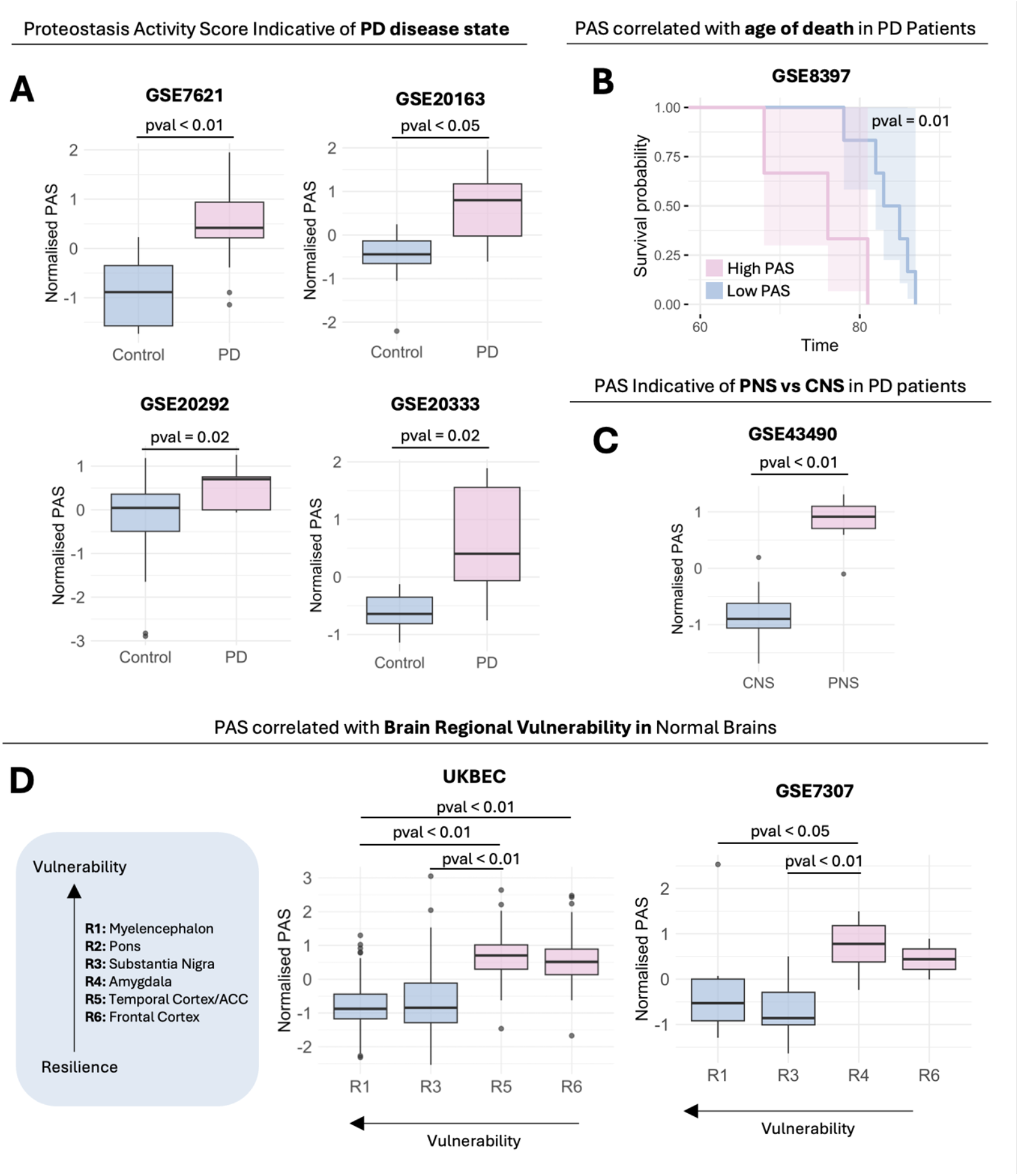
The proteostasis activity score (PAS) is indicative of PD disease states, age of death, and brain regional vulnerability. **(A)** PAS in PD brains are higher than control brains. **(B)** Higher PAS is associated with younger age of death in PD patients. **(C)** In PD patients, regions that exhibit earlier α-Syn aggregation in disease (vagus nerve, peripheral nervous system) have a higher PAS than in the central nervous system. **(D)** We extended the analysis to healthy brains, finding that the PAS are indicative of regional vulnerability in the brain, with more vulnerable regions having a lower baseline PAS than regions that are more resilient. The Wilcoxon test was used for calculating statistical significance.

### Target identification for PD drug discovery

To propose potential protein targets for PD drug discovery, we hypothesized that proteins that exert larger influence on the α-Syn PN are more likely to be useful targets. To investigate this possibility, we quantified two network influence scores – heat diffusion and personalised page rank (PPR). Heat diffusion quantifies the signal of a protein spread across the network to understand the its influence over signal propagation in the network itself. The proteins are plotted on a 2-axis graph in **Figure 3A**. Proteins with high heat diffusion and high PPR are likely to be master regulators, as they are central hubs and efficiently propagate signals through the network. Proteins with high heat diffusion and lower PPR efficiently, although not major hubs, propagate signals locally and likely modulate specialized functions within the network. Proteins with high PPR but lower heat diffusion are well connected, but they are less efficient at propagating signals through the network. We identified 55% of the proteins making up our α-Syn PN as target proteins for PD in the Open Targets database **(Figure 3B)**. Of these proteins, signal modulators and essential proteins with higher heat diffusion and signal modulation capabilities tended to have relatively higher priority scores for PD as per Open Targets **(Figure 3C)**. Notably, one third of these essential/signal modulators are proteins that have yet to be reported to be targets for PD **(Figure 3D)**. These proteins are PIK3CD, KIAA0319, HSPA2, TBC1D4, IMP4, and VPS37C. Experimental inhibition of the target proteins beneficially modulate α-Syn PN activity as quantified by the PAS **(Figure 3E)**.

**Figure 3:**
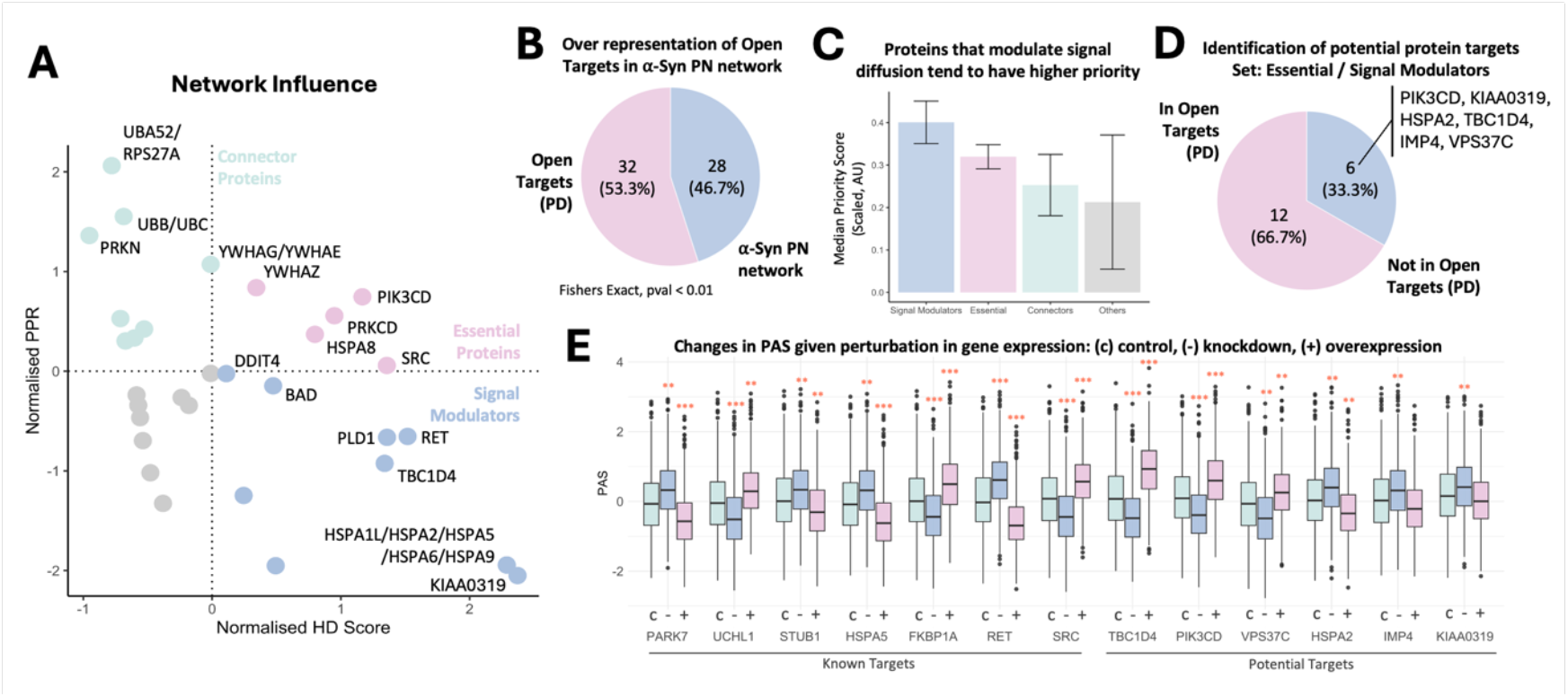
Quantitative network influence facilitates the identification and prioritization of potential target proteins. **(A)** The network influence of each protein was quantified using the heat diffusion and personalised page rank (PPR) algorithm. Proteins with high heat diffusion and high PPR are likely to be master regulators as they are central hubs and efficiently propagate signals through the network. Proteins with high heat diffusion and lower PPR efficiently, although not major hubs, propagate signals locally and likely modulate specialized functions within the network. Proteins with high PPR but lower heat diffusion are well connected but are less efficient at propagating signals through the network. **(B)** Of the 60 proteins in the α-Syn PN, 33 (55%) have been identified to be target proteins for PD in the Open Targets database. **(C)** Signal modulators and essential proteins, proteins with higher heat diffusion and signal modulation capabilities, tend to have relatively higher priority scores for PD as per Open Targets. **(D)** Six of 18 essential/signal modulators are novel proteins that have yet to be reported to be target proteins for PD. The 6 proteins are PIK3CD, KIAA0319, HSPA2, TBC1D4, IMP4, and VPS37C. **(E)** Simulated inhibition or activation of the target proteins through a digital twin approach (see **Methods**) beneficially modulate the α-Syn PN activity as quantified by PAS. Effect size of gene-wise expression perturbations were calculated using Cohen’s D: (***) Cohen’s D > 0.5 and (**) Cohen’s D > 0.3.

To study the modulatory effects potential targets have on the activity of the α-Syn PN, we created a digital twin of brain cells in the *Substantia nigra* to simulate the knockdown and overexpression of selected known targets and potential targets in order to prioritize experiments (see **Methods**). By benchmarking the simulation to known targets **(Figure 3E)**, we found that downregulation of α-Syn aggregation-attenuating PARK7 resulted in a relatively higher PAS score indicating a shift of the α-Syn PN activity toward promoting α-Syn aggregation. The overexpression of α-Syn aggregation-promoting UCHL1 also increased the aggregation-promoting activity of the network, while taking a more inhibitory slant upon downregulation. Furthermore, α-Syn aggregation-inhibiting STUB1 and RET shifted the activity of the network towards inhibiting α-Syn aggregation upon upregulation while causing a slant towards aggregation-promoting activity upon their downregulation. Our top 3 potential targets based on their effect size on the α-Syn PN given expression perturbations are TBC1D4, PIK3CD, and VPS37C **(Figure 3E)**. All 3 targets were upregulated relative to *SNCA* in PD conditions and are found to pivot the activity of the network toward promoting α-Syn aggregation upon overexpression. In contrast, downregulating these targets shifted the network closer toward α-Syn inhibition. These observations suggests that inhibition of TBC1D4, PIK3CD, and VPS37C may potentially help manage the shift of the α-Syn PN toward promoting aggregation.

### Drug prioritization for repurposing

In this section, we used the a-Syn PN to define a metric to help propose and prioritize drugs for PD. Towards this end, we first selected cell models that best capture the shift in the α-Syn PN towards aggregation. We carried out a comparison of 1206 cell lines listed in the Human Protein Atlas based on their relative PAS **(Figure 4A)**. Cell lines with higher PAS indicating α-Syn PN activities slanted toward α-Syn aggregation promotion were given higher priority. From here, two model cell lines available in LINCS (a large-scale drug testing dataset) were identified: Jurkat (rank 30) and HEK293 (rank 48). Corroboration with literature suggests that both HEK293 (38-41) and Jurkat cells (42, 43) may be relevant for PD-related studies. We then obtained differential gene expression data for both cell lines treated with various small molecules (968 drugs tested in HEK293 cells and 1056 drugs tested in Jurkat cells) compared against matched cells treated with DMSO. Changes in PAS (ΔPAS) upon treatment were quantified for each drug at each concentration tested (treatment for 24 h). For each drug, we then calculated the correlation of increasing levels of drug concentrations with ΔPAS in both HEK293 **(Figure 4B)** and Jurkat **(Figure 4C)** cells. 813 drugs were tested in both cell lines, with 435 of those drugs (36%) exhibiting consistent effects on the PAS in both cells **(Figure 4D)**. We highlighted 28 inhibitors of α-Syn PN that significantly shift the α-Syn PN activity toward inhibiting α-Syn aggregation. To facilitate drug repurposing for PD, we prioritized the 28 inhibitory drugs according to their average absolute correlation score across HEK293 and Jurkat cells **(Figure 4E)**.

**Figure 4.**
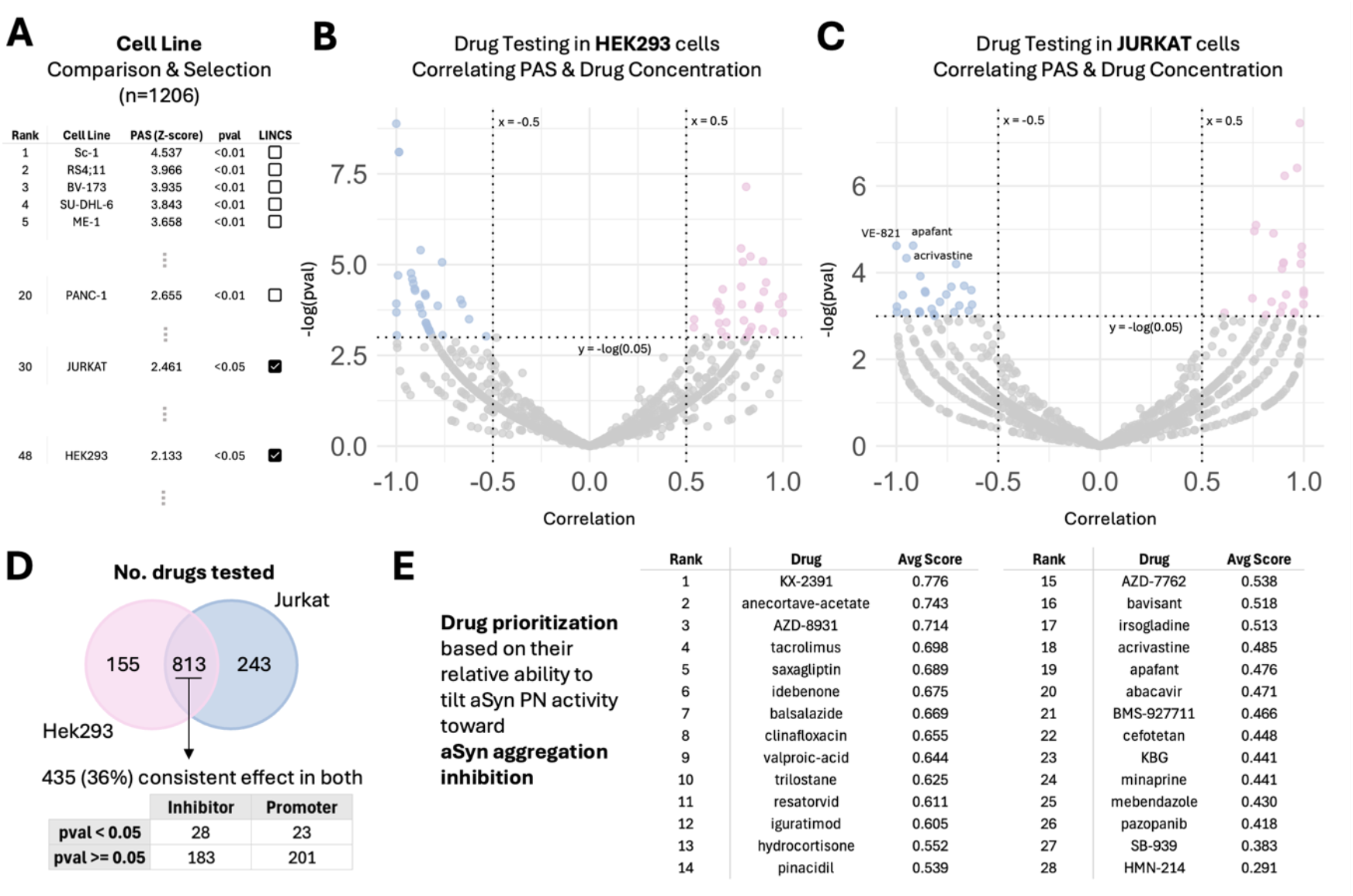
The quantification of changes in PAS given drug treatment in HEK293 and Jurkat cells enables the identification and priorization of repurposed drugs modulating α-Syn PN toward inhibition of α-Syn aggregation. **(A)** To select the best model for drug testing, we compared the α-Syn PN activity (quantified by the PAS) for 1206 cell lines in the Human Protein Atlas (HPA). We ranked the cell lines based on their relative PAS activity, prioritizing cell lines that have a α-Syn PN activities slanted toward α-Syn aggregation promotion (higher PAS). **(B**,**C)** Based on this ranking, we identified 2 cell lines: Jurkat (rank 30) and HEK293 (rank 48) that are available within the LINCS drug testing dataset. For both cell lines, we obtained differential gene expression data for the cells treated with various small molecules compared against matched cells treated with DMSO. Using these data, we calculated the change in PAS (ΔPAS) upon treatment with different small molecules at increasing concentrations for 24 h. For each drug, we calculated the correlation of increasing levels of drug concentrations with ΔPAS in both HEK293 (B) and Jurkat (C) cells. **(D)** Of the 968 drugs tested in HEK293 cells and 1056 drugs tested in Jurkat cells, 813 drugs were tested in both, with 435 of the 813 common drugs (36%) exhibiting consistent effects on the PAS in both cells. Notably, 28 inhibitors of α-Syn PN (drugs with negative correlation between drug concentration and ΔPAS in both cell lines) significantly (p-value <0.05 in at least one of the cell lines) shift the α-Syn PN activity toward inhibiting α-Syn aggregation. **(E)** To facilitate drug repurposing for PD, we prioritized the 28 inhibitory drugs according to their average absolute correlation score across HEK293 and Jurkat cells.

This approach enables the prediction of repurposed drugs potentially relevant for PD. About one third of the drugs that we identified have already been studied and reported to be beneficial in PD models. Some examples are: (i) Tacrolimus, originally designed for preventing post-transplant organ rejection, was reported to lead to improvements in the functional features of dopaminergic neurons and behavioural phenotypes affected in PD at low doses in mice (44); (ii) Saxagliptin, originally an anti-diabetic drug, was reported to significantly improve motor performance, muscle coordination and correct akinesia in rat PD models (45), and proposed as a novel approach toward the management of PD (45); (iii) Idebenone, originally prescribed for dementia, was reported to improve motor dysfunction, learning and memory (46), as well as, reduce neuroinflammation (47) in PD mice models; (iv) Hydrocortisone, usually prescribed for to relieve inflammation, prevents dopaminergic cell in a PD cell line model (48); (v) Pinacidil, usually used to control blood pressure, was found to exhibit anti-inflammatory effects in an in vitro PD microglia model (49); (vi) Bavisant, originally developed to treat ADHD, has completed phase 2 trials to treat PD-related symptoms (ClinicalTrials.gov, identifier NCT03194217); (vii) Minaprine was identified as a PD drug in ChEBI (https://www.ebi.ac.uk/chebi/searchId.do?chebiId=51038), and (viii) Pazopanib, the first FDA-approved for treating advanced renal cell carcinoma, which has anti-inflammatory and neuroprotective effects in dopaminergic neurons in mouse models (50).

## Discussion

Aberrant proteostasis is likely to contribute to PD, as reduced degradation of α-Syn results in the accumulation of aggregates associated with a toxic gain-of-function (24-26, 31-36, 51-54). In this study, we investigated the links between PD and the progressive impairment of the proteostasis system that regulates α-Syn. This approach followed previous studies that related neurodegenerative conditions with the a dysregulation of proteostasis (18-23).

We generated a representation of the α-Syn PN using proteomic and transcriptomic data. Components of the α-Syn PN were then cross-checked with literature and across the Braak staging. To illustrate the application of the α-Syn PN, we used it to study of disease states in PD patients, age-of-death, and brain vulnerability toward α-Syn aggregation in PD. We further showed how the α-Syn PN can be applied toward translational applications such as target identification and drug repurposing.

To facilitate the analysis, we defined a score (PAS) that quantitatively describe the α-Syn PN activity and its effects in modulating α-Syn aggregation. We found that the PAS helps represent disease states, whereby higher PAS, which indicate promotion of α-Syn aggregation, is associated with PD disease states compared to controls, a younger age-of-death amongst PD patients, and in the PNS system that is susceptible earlier in PD compared to the CNS. We also found that in control brain samples, brain regions that are more vulnerable to PD tend to have lower basal PAS than relatively more resistant brain regions. This result aligns with previous studies finding lower basal *SNCA* levels in more vulnerable brain regions compared to more resilient regions (55). These findings suggest that regions with a higher susceptibility toward α-Syn aggregation (lower PAS scores and lower basal levels of *SNCA*) present disease features earlier than regions with higher resilience toward α-Syn aggregation (higher PAS scores and higher basal levels of *SNCA*). Taken together, these observations suggest that before the onset of the disease, a high proteostasis activity is protective, while after the onset of the disease, a high proteostasis activity emerges a consequence of the disease itself.

We then investigated whether modeling a digital twin of the α-Syn PN for a brain cell in the *Substantia nigra* can help experimental efforts in target identification. We showed that the study of information flow across the a-Syn PN leads to the identification of relevant PD targets. To propose new targets, the digital twin captures protein-protein dependencies of the α-Syn PN. This approach offers two main benefits. First, the digital twin allows for the simulation of complex processes to enable the study of network responses to perturbations such as knockdown, overexpression, and can be extended to drug perturbations. By doing so, one can carry out a first screen, and then filter the targets to be taken forward into experimental stages - thus potentially saving time, effort and financial costs. Second, the digital twin of cells that are hard to access, such as those in brain tissues, which can most often only be sampled post-mortem, enables the study of complex biological responses in these environments that complements drug testing cell lines. While we acknowledge that digital twin approaches of the type adopted in this work are still in their early stages, we highlight their potential in enabling simulated drug testing directly in disease-relevant tissues especially for hard-to-access tissues.

In a second case study, we analysed drug repurposing based on measuring changes of the PAS in response to drug treatment. This analysis, which was done in HEK293 and Jurkat cells, yielded 28 drugs predicted to shift the α-Syn PN activity towards the inhibition of α-Syn aggregation. As part of the analysis, we compared 1026 cell lines listed in the Human Protein Atlas based on their relative PAS. Intriguingly, we found that the top 10 cell lines in terms of PAS were lymphoblasts/leukemia cell lines. Consistent with these findings, α-Syn has been reported to be differentially expressed in blasts of leukemia cells and has been proposed as a diagnostic marker (56, 57). In addition, a role for α-Syn has been previously proposed in the function of lymphocytes (58). Further studies will be needed to investigate the relevance of this association.

In conclusion, we hope that the definition of the α-Syn PN reported in this study could serve as a reference for further studies of PD disease mechanisms and potential diagnostic and therapeutic interventions.

## Methods

### PN data and their functional interactions

A comprehensive list of PN was obtained from the Proteostasis Consortium (59, 60)(https://www.proteostasisconsortium.com/). Functional pairwise interactions (version 2021) were downloaded from the Reactome database (https://reactome.org/). All predicted interactions were filtered out from this set of pairwise interactions obtained from Reactome.

### Identification of the first-degree PN regulators of α-Syn

To identify the PN proteins that directly exert a functional interaction on α-Syn, we filtered the pairwise interactions for any inward functional interactions from any protein within the Proteostasis Consortium list of PN proteins with α-Syn. The unique list of proteins obtained from the one-directional edgelist resulted in a set of first-degree α-Syn interactors (also referred to as primary α-Syn PN) visualised in **Figure 1**. To ensure relevance of the first-degree PN proteins found, we carried out a literature search for evidence supporting their regulatory roles in α-Syn aggregation **(Table S1)**. Based on the reports found in literature, we further classified them into promoters and ‘attenuators of α-Syn aggregation.

### PD microarray datasets and consistently perturbed genes relative to *SNCA*

To identify a consensus set of genes whose expression in relation to *SNCA* is altered in PD compared to controls, we downloaded and analysed 6 PD microarray datasets from the NCBI Gene Expression Omnibus (GEO) (www.ncbi.nlm.nih.gov/geo/): GSE8397 (61), GSE7621 (62), GSE20163 (63), GSE20292 (63), GSE20333, and GSE43490 (64). These datasets were selected as they had minimally 5 control and 5 disease samples from the *Substantia nigra*. All differential analysis between disease and control samples were done separately for each dataset before the results were pulled together for further comparison. All expression values were log2-transformed. To calculate the relative expression of each gene to *SNCA* (for evaluating balance of proteins with respect to α-Syn), we normalise all genes to *SNCA* expression by taking log2([gene]/[*SNCA*). Statistical differences between the relative expressions (gene:*SNCA*) between groups were evaluated using the two-sided Wilcoxon test with Bonferroni correction for false-discovery detection. A cutoff of absolute fold-change > 0.3 and FDR < 0.05 was used. Based on these thresholds, 108 genes were found to be consistently altered (directionality) in at least half of the datasets analysed **(Table S2)**.

### Incorporating consensus genes into the α-Syn PN

Seeking to incorporate the relevant subset of the 108 consensus genes into the α-Syn PN, we looked for pairwise functional interactions between each of the 108 genes with the existing primary first-degree PN interactors of α-Syn. From here, 21 genes were found to have functional interactions with the first-degree primary α-Syn PN established earlier and were included into the final α-Syn PN **(Table S3)**. These 20 genes were determined to be upstream genes potentially capable of regulating the α-Syn PN and were hence added into the α-Syn PN. Inward functional interactions between each of the 108 genes and α-Syn itself were also searched for. However, none of the 108 genes were found to exert inward functional activities on α-Syn within this dataset.

### Validating the 20 upstream genes within the α-Syn PN

To validate our consensus gene set, we downloaded an additional microarray dataset GSE49036 (65) which had information on Braak staging for comparison. Similar to before, all expression values were log2-transformed, normalised to *SNCA*, and visualised in a boxplot by Braak staging.

### Proteostasis activity score (PAS)

We defined the proteostasis activation score in a similar way to the subnetwork expression metric used in the context of protein-interaction networks (66). The metric over the network of size K is defined as

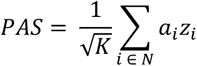

where *z*_*i*_ denotes the Z-score normalised expression profile of gene *i* across the all genes and *a*_*i*_ is the pathway activation sign with *a*_*i*_ = 1 if activating and *a*_*i*_ = -1 if inactivating/inhibitory.

### Influence scores of primary α-Syn PN components

To quantify the relative importance of each protein within the α-Syn PN, the influence of each protein score was used as a proxy for its importance within the network. The heat diffusion (67) and personalized page rank (68) algorithms were applied to all proteins within the α-Syn PN network to estimate their influence. The influence scores are available in **Table S4**.

### Non-disease multiple brain region datasets

Two human datasets including non-disease samples from various brain regions were used to study the association of our genes with regional vulnerability to PD: UKBEC (37) and GSE7307. Six Braak-stage related regions were delineated R1-R6 (of increasing resilience/decreasing vulnerability) as previously reported (55): R1 – myelencephalon, R2 – pons, R3 – *Substantia nigra*, R4 – amygdala, R5 – temporal cortex/ACC, and R6 – frontal cortex. Samples from brain regions R1, R3, R5, R6 were available within UKBEC and downloaded for analysis. The biomaRt package was used to map the Affymetrix probe IDs from the UKBEC dataset to gene symbols in R. Where multiple probes mapped to the same gene, the probe with the highest variance across samples was chosen. All expression values were expressed on the log2-scale and normalised to *SNCA* expression sample-wise as described earlier. Relative expression values were compared and visualised by brain regions.

### Digital twins of the α-Syn PN in brain cells in the *Substantia nigra*

Aiming to accelerate the discovery of PD-related mechanisms and drug testing, we created a virtual replicate of our α-Syn PN. For this purpose, we built a digital twin of the α-Syn PN for brain cells in the *Substantia nigra*. Our goal was to enable the simulation of complex biological processes to gain insights into disease mechanisms and to evaluate potential treatments and their effects on biological networks. By forecasting how individual components of the network would change given perturbations, a digital twin could be useful in assisting in translational applications such as target identification and drug discovery. To build the digital twin, single-cell transcriptomic data from *Substantia nigra* brain samples were obtained from the human single cell atlas of the *Substantia nigra* (GSE140231) (69). Single-cell gene expression data were processed using the Seurat package (70) in R. The Z-score normalisation was applied to normalise gene expressions across all genes in each cell. Z-scores for the α-Syn PN components were extracted and used to learn the dependencies between components of our α-Syn PN via their partial correlations by fitting graphical Gaussian models (GGMs) using the GeneNet package in R (https://cran.r-project.org/web/packages/GeneNet/index.html). This method was previously shown to outperform other methods in predicting network structure and dependencies from gene expression data (71, 72). The resultant dependencies were filtered using p-value <0 .05 and used to fit a linear model allowing the prediction of expression changes for each gene in the network given perturbations to a query gene. All single cells within the dataset that passed quality control (QC) were used for training. Since this procedure did not differentiate for cell type, our digital twin represents an average brain cell of the *Substantia nigra*. We note that in future studies, digital twins of specific cell types could be generated using the approach used here, subject to the availability of appropriate datasets.

### Gene expression perturbation simulation

To simulate knockdown and overexpression experiments, we modulated the expression value of each query gene by the following – knockdown: 0.06, 0.12, 0.25, 0.5, 0.75 times the basal level, and overexpression: 1.25, 1.5, 1.75, 2, 4, 8 times the basal level in each cell. We then predicted new gene expression levels for the other genes within the network. Using the new expression levels, we calculated an updated PAS for each cell. For visualisation in **Figure 3**, gene downregulation samples were gene expression levels given perturbation of a query gene at 0.12 times the basal level, while overexpression was gene expression levels given perturbation of a query gene at 8 times the basal level in each cell. Detailed simulated data at varying concentrations of downregulation and overexpression for known benchmark targets and our top 3 potential targets are reported in **Figure S3**.

### Comparison and selection of cell line models for drug testing

nTPM gene expression values for 1026 cell lines were obtained from the Human Protein Atlas (https://www.proteinatlas.org/). To obtain relative gene expression levels, we calculated the ratio of all genes to *SNCA*. Next, Z-score normalisation was applied to the relative gene expression ratios. The gene expression levels for genes within the α-Syn PN were extracted for each cell line and a corresponding PAS was calculated. Finally, a Z-score normalisation was applied to normalise the PAS across all cell lines. A p-value was also calculated, with p-value <0.05 taken as the cutoff for significance. The cell lines were ranked in accordance from highest PAS to lowest PAS, prioritizing cell lines with relatively more α-Syn promoting PNs.

### LINCS dataset

CMAP LINCS 2020 level 5 data (level5_beta_trt_cp_n720216×12328.gctx) was downloaded from clueio (https://clue.io/data/CMap2020#LINCS2020). Only data from HEK293 and Jurkat cells treated with small molecules were extracted for use in this study. For each condition (small molecule treatment at a given concentration), the change in PAS (small molecule treatment vs DMSO/control) was calculated to understand the effects of the small molecule on the α-Syn PN. The change in PAS (ΔPAS) was defined as

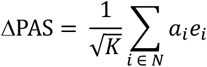

where *e*_*i*_ denotes the differential relative expression (differential expression of genei:*SNCA*) of a gene *i* while *a*_*i*_ is the pathway activation sign with *a*_*i*_ = 1 if activating and *a*_*i*_ = -1 if inactivating/inhibitory for the α-Syn PN sized K.

For each of HEK293 and Jurkat cell models, the correlation between the increasing concentrations of each drug (log scale) and the ΔPAS were calculated. A p-value <0.05 and absolute correlation score of 0.5 was used as cutoff. Drugs with negative correlation scores were classified as inhibitors, shifting the activity of the α-Syn PN toward preventing α-Syn aggregation. In contrast, drugs with positive correlation scores were calculated as promoters, shifting the activity of the α-Syn PN toward promoting α-Syn aggregation.

### Drug prioritization

For the use of re-purposing drugs for PD, we prioritized drugs with inhibitory effects on α-Syn aggregation via the α-Syn PN. Of the 813 drugs tested in both cell lines, 28 drugs were found to consistently exhibit inhibitory effects on α-Syn aggregation via the α-Syn PN. These drugs were ranked based on their average absolute correlation scores in each cell line.

