## Supplementary Information for "The α-Synuclein Proteostasis Network and its Translational Applications in Parkinson’s disease"

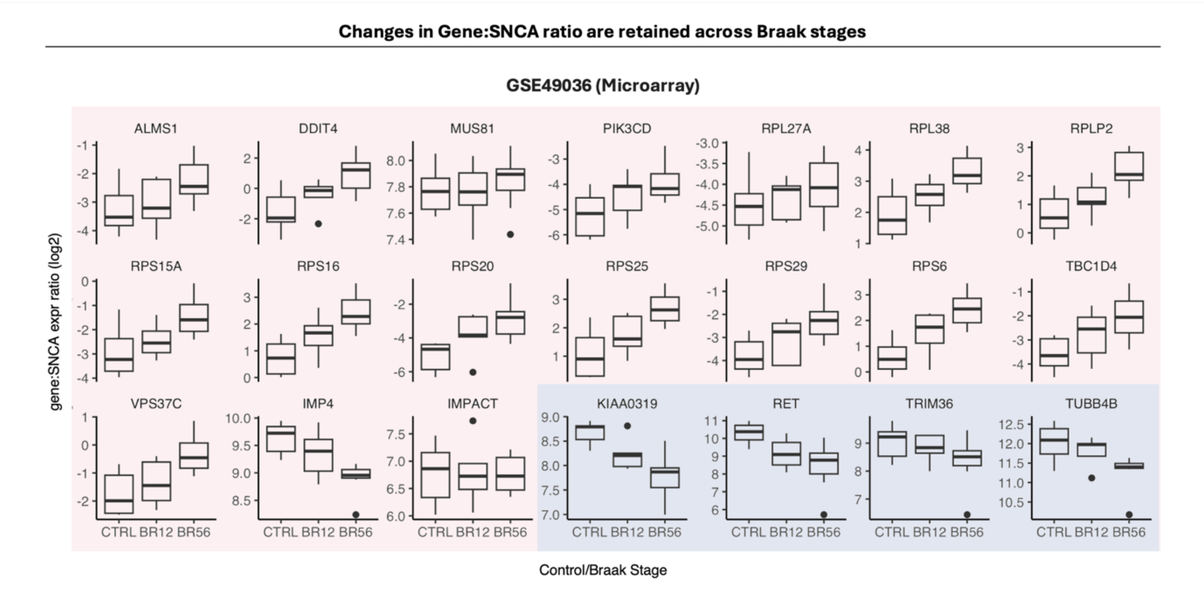


**Figure S1. Validation of the perturbation trends of the proteins in the a-Syn PN across Braak stages in PD patients.** 108 genes were found to be consistently perturbed relative to *SNCA* in PD brains in majority of the 6 PD patient datasets analysed as described in **Methods**. Of these 108 genes, 21 were found to encode proteins that have functional interactions with the first-degree primary α-Syn PN **(Table S1)**. To validate their perturbation patterns in PD, we analysed GSE49036 which is a microarray dataset consisting of *Substantia nigra* samples from PD brains of different Braak stages and find that the same trend of perturbation is retained **(Table S2)**.

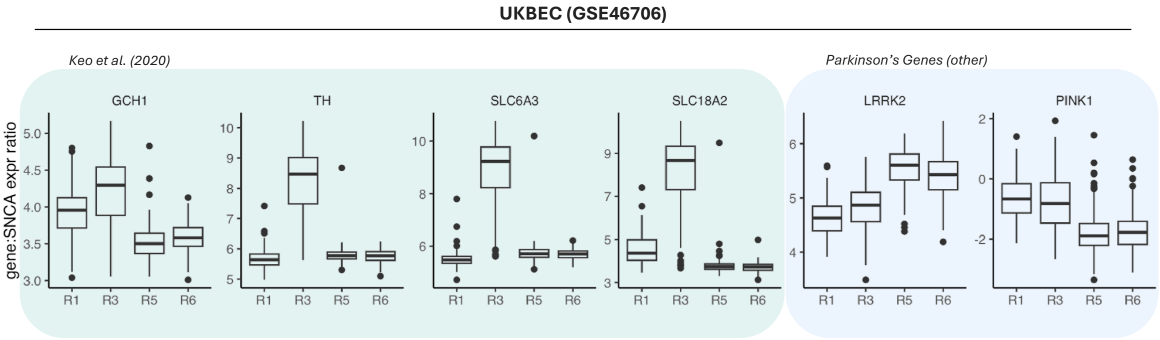


**Figure S2: Benchmarking of other PD-associated genes.** Genes identified via differential expression[24] may not always be indicative when considered in relation to α-Syn.

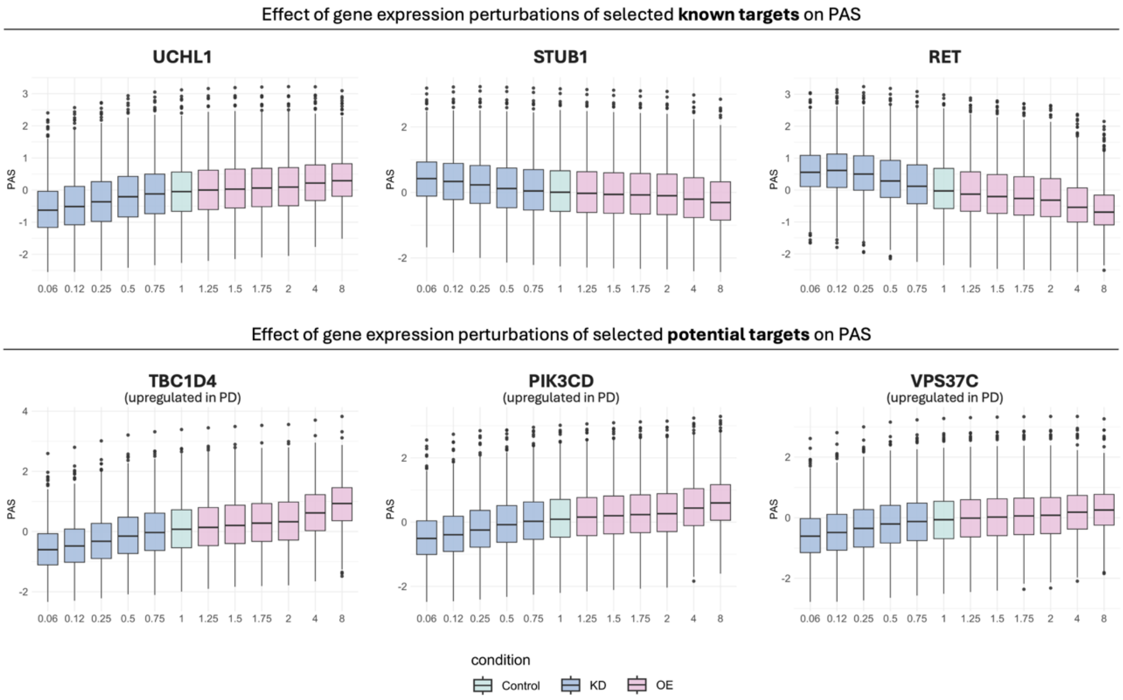


**Figure S3. Effect of gene expression pertubations of selected known PD targets and potential PD targets on the activity of the α-Syn PN.** To study the modulatory effects potential targets have on the activity of the α-Syn PN, we simulated the knockdown and overexpression of selected known targets and potential targets to mimic corresponding experiments in cells. Details of the simulations are described in **Methods**. Benchmarking our simulations to known targets, we found that overexpression of α-Syn aggregation-promoting UCHL1 increased the aggregation-promoting activity of the network upon upregulation and took a more inhibitory slant upon downregulation. α-Syn aggregation-inhibiting STUB1 and RET shifted the activity of the network towards inhibiting α-Syn aggregation upon upregulation while causing a slant towards aggregation-promoting activity upon their downregulation. Our top 3 potential targets based on their effect size on the α-Syn PN given expression perturbations are TBC1D4, PIK3CD, and VPS37C. All 3 targets are upregulated relative to *SNCA* in PD conditions and are found to pivot the activity of the network towards promoting α-Syn aggregation upon overexpression. In contrast, downregulating these targets shifted the network closer toward α-Syn inhibition. These results suggest that inhibition of TBC1D4, PIK3CD, and VPS37C may potentially be leveraged for managing the shift of the α-Syn PN towards promoting aggregation.
